# Individual prediction tendencies do not generalise across modalities

**DOI:** 10.1101/2023.02.02.526758

**Authors:** Juliane Schubert, Nina Suess, Nathan Weisz

## Abstract

Predictive processing theories, which model the brain as a “prediction machine”, explain a wide range of cognitive functions, including learning, perception and action. Furthermore, it is increasingly accepted that aberrant prediction tendencies play a crucial role in psychiatric disorders. Given this explanatory value for clinical psychiatry, prediction tendencies are often implicitly conceptualised as individual traits or as tendencies that generalise across situations. As this has not yet explicitly been shown, in the current study, we quantify to what extent the individual tendency to anticipate sensory features of high probability generalises across modalities. Using magnetoencephalography (MEG), we recorded brain activity while participants were presented a sequence of four different (either visual or auditory) stimuli, which changed according to predefined transitional probabilities of two entropy levels: ordered vs. random. Our results show that, on a group-level, under conditions of low entropy, stimulus features of high probability are preactivated in the auditory but not in the visual modality. Crucially, the magnitude of the individual tendency to predict sensory events seems not to correlate between the two modalities. Furthermore, reliability statistics indicate poor internal consistency, suggesting that the measures from the different modalities are unlikely to reflect a single, common cognitive process. In sum, our findings suggest that quantification and interpretation of individual prediction tendencies cannot be generalised across modalities.

## INTRODUCTION

In our complex sensory environments, the amount, as well as the level of ambiguity, of information entering our senses requires a system that can filter and integrate information efficiently. As such a system, our brain has to rely fully on its sensory input to infer the true states of objects that cause such input. This inference results in the formation of a so-called internal model, which exists so that predictions can be drawn from it. Even though the assumed implementations of these predictions vary across different “predictive brain” theories (Friston, 2010; Knill & Pouget, 2004; Yon et al., 2019), most agree on the importance of predictions for perception in general. Bottom-up-driven sensory representations in the brain inevitably lag behind the events that caused them, and therefore compensatory mechanisms such as the prediction of future events are highly beneficial for our everyday lives (e.g. to correctly allocate moving objects). Indeed, such anticipation and pre-activation of sensory input has been found for visual perception (Hogendoorn & Burkitt, 2018), as well as for language processing (Dikker & Pylkkänen, 2013). Furthermore, we recently showed that “prediction tendency”, which we defined as the tendency to anticipate auditory features of high probability before their occurrence, contributes to explaining individual differences in speech tracking (Schubert et al., 2023).

Complementary to our results, various research has pointed out that individuals seem to vary in the extent to which they rely on top-down priors compared to bottom-up sensory signals for perception. Crucially, these differences in handling predictions or in overall “prediction tendency” have been associated with clinical psychological conditions such as autism (Sinha et al., 2014), schizophrenia (Sterzer et al., 2018), depression (Barrett et al., 2016), PTSD (Kube et al., 2020) and tinnitus (Partyka et al., 2019; Sedley et al., 2016). Taking its explanatory value to clinical psychiatry into account, prediction tendency is often implicitly conceptualised as an individual trait or as a tendency that generalises across situations. Indeed, in line with predictive brain theories for psychosis (Sterzer et al., 2018), it has been found that strong predictive tendencies can promote phantom perception in the auditory modality (Powers et al., 2017). Together with our findings, which state that the aforementioned internal mechanisms are related to differences in speech tracking (Schubert et al., 2023), this strongly suggests that individual prediction tendencies are generalisable across different listening situations. However, the extent to which they generalise across different modalities remains unclear.

Furthermore, there are considerable differences between different modalities (e.g. auditory vs. visual) in the predictability of sensory information and, therefore, also in the way predictions should best be applied. Visual information (e.g. a single object or a whole scene) naturally unfolds across space, and sensory inference is often drawn from spatially surrounding information. In audition, however, this is not necessarily the case. Considering speech and music, for example, whose predictability mainly unfolds over time, which seems to be a core characteristic of auditory objects. It follows that, in the auditory domain, two types of predictions are usually distinguished, carrying spectral (*what is going to happen*) or temporal (*when is it going to happen*) information (Auksztulewicz et al., 2018; Wollman & Morillon, 2018). In the visual modality, on the other hand, the focus is often on object-based (*what is it*) and spatial (*where is it*) predictions, based on dual-stream theories of vision (e.g. Rao & Ballard, 2005). Nevertheless, a correct anticipation of sensory events should include all of these aspects (*what, when and where*) in both modalities, especially since predictions that generalise and complement across modalities are highly beneficial in natural settings (integrating, for example, lip movements and acoustic speech; Crosse et al., 2016; Sumby & Pollack, 1954). If “prediction tendency” can indeed be conceptualized as an individual trait, it should be similar across situations, as well as across modalities. Complementing correlation estimates, standard measures of internal consistency should help in evaluating whether measures from the auditory and visual modality reflect a putatively common construct. In such a case, internal consistency measures should be sufficiently high (at least >= .7; Tavakol & Dennick, 2011). Inherent differences between auditory and visual perception, however, emphasise the importance of comparative research in the field of predictive processing.

In the current study, we used magnetoencephalographic (MEG) data in visual and auditory versions of an entropy modulation paradigm (see also Demarchi et al., 2019; Schubert et al, 2023) to capture individual prediction tendencies for both modalities. In this paradigm, participants were presented with a sequence of four different (either visual or auditory) stimuli, which changed according to predefined transitional probabilities of two entropy levels: ordered vs. random. In order to optimise comparability, we kept constant spatial predictability (visual stimuli were presented at screen center and auditory stimuli were presented binaurally at phantom center), as well as temporal predictability (with a fixed stimulation rate of 3 Hz), modulating only one stimulus feature in each modality. In the visual version, participants were presented with a sequence of gabor patches that changed in orientation, whereas in the auditory version, participants listened to pure tones of different frequencies. For both modalities, individual prediction tendency was defined as the tendency to anticipate and pre-activate stimulus features of high probability.

## METHODS

### Subjects

In total, 35 subjects (16 male, mean age = 32.5, range = 19 - 57) were recruited to participate in the experiment. All participants reported normal hearing and had normal, or corrected to normal, vision. They gave written, informed consent and reported that they had no previous neurological or psychiatric disorders. The experimental procedure was approved by the ethics committee of the University of Salzburg and was carried out in accordance with the declaration of Helsinki. All participants received either a reimbursement of 10 € per hour or course credits for their participation.

### Experimental Procedure

Before the start of the experiment, participants’ individual head shapes were assessed using cardinal head points (nasion and pre-auricular points), digitised with a Polhemus Fastrak Digitiser (Polhemus), as well as around 300 points on the scalp. For every participant, MEG sessions started with a 5-minute resting-state recording, after which individual hearing threshold was determined, using a pure tone of 1043 Hz. This was followed by 4 experimental blocks (2 blocks per modality) of an entropy modulation paradigm. Participants started with either the auditory or visual blocks (chosen randomly for each participant). In the auditory paradigm (see also Schubert et al., 2023), participants passively listened to sequences of four different pure tones (1: 440 Hz, 2: 587 Hz, 3: 782 Hz, 4: 1043 Hz) while watching a landscape movie (LoungeV Films, 2017). All tones were presented binaurally at equal volume for the left and right ears (i.e. at phantom center) at 40 db above the individual hearing threshold. In the visual paradigm, a gabor patch (spatial frequency: 0.01 cycles/pixel, sigma: 60 pixels, phase: 90°) was presented at four perceptually different orientations (1: 0°, 2: 45°,3: 90°, 4: 135°) in the center of the screen while participants listened to podcasts from “Bayern 2 - radioWissen”: “Kakao - Das braune Gold” (Cocoa - the brown gold) and “Zimt - Die Würze des Lebens” (Cinnamon - the spice of life). There was no task, and participants were just instructed to relax and to move as little as possible. To ensure temporal predictability, all stimuli were presented at a fixed stimulation rate of 3 Hz, and each stimulus presentation (i.e. pure tone or gabor patch) lasted for 100ms. One block consisted of a sequence of 2800 trials, totalling 5600 trials per modality. Transitional probabilities between stimuli (1, 2, 3, 4) were determined by two different entropy levels (ordered vs. random see **Fig. 1A**) . Entropy changed pseudorandomly within each block every 700 trials. While in an “ordered” context, certain transitions (hereinafter referred to as forward transitions, i.e. 1→2, 2→3, 3→4, 4→1) were to be expected, with a high probability of 75%, self repetitions (e.g., 1→1, 2→2,…) were rather unlikely, with a probability of 25%. However, in a “random” context, all possible transitions (including forward transitions and self repetitions) were equally likely, with a probability of 25% (see **Fig. 1A** and see also Schubert et al., 2023). Furthermore, a pseudorandom 10% of the trials (i.e. ∼ 70 trials per stimulus, per entropy, per modality) were omitted (as in Demarchi et al., 2019). Between each of the four blocks (∼15 min each), there was a short break. In total, the experiment lasted approximately 2h per participant (including MEG preparation time). The experiment was coded and conducted with the Psychtoolbox-3 (Brainard, 1997; Kleiner et al., 2007), with an additional class-based library (‘Objective Psychophysics Toolbox’, o_ptb) on top of it (Hartmann & Weisz, 2020).

**Fig. 1:**
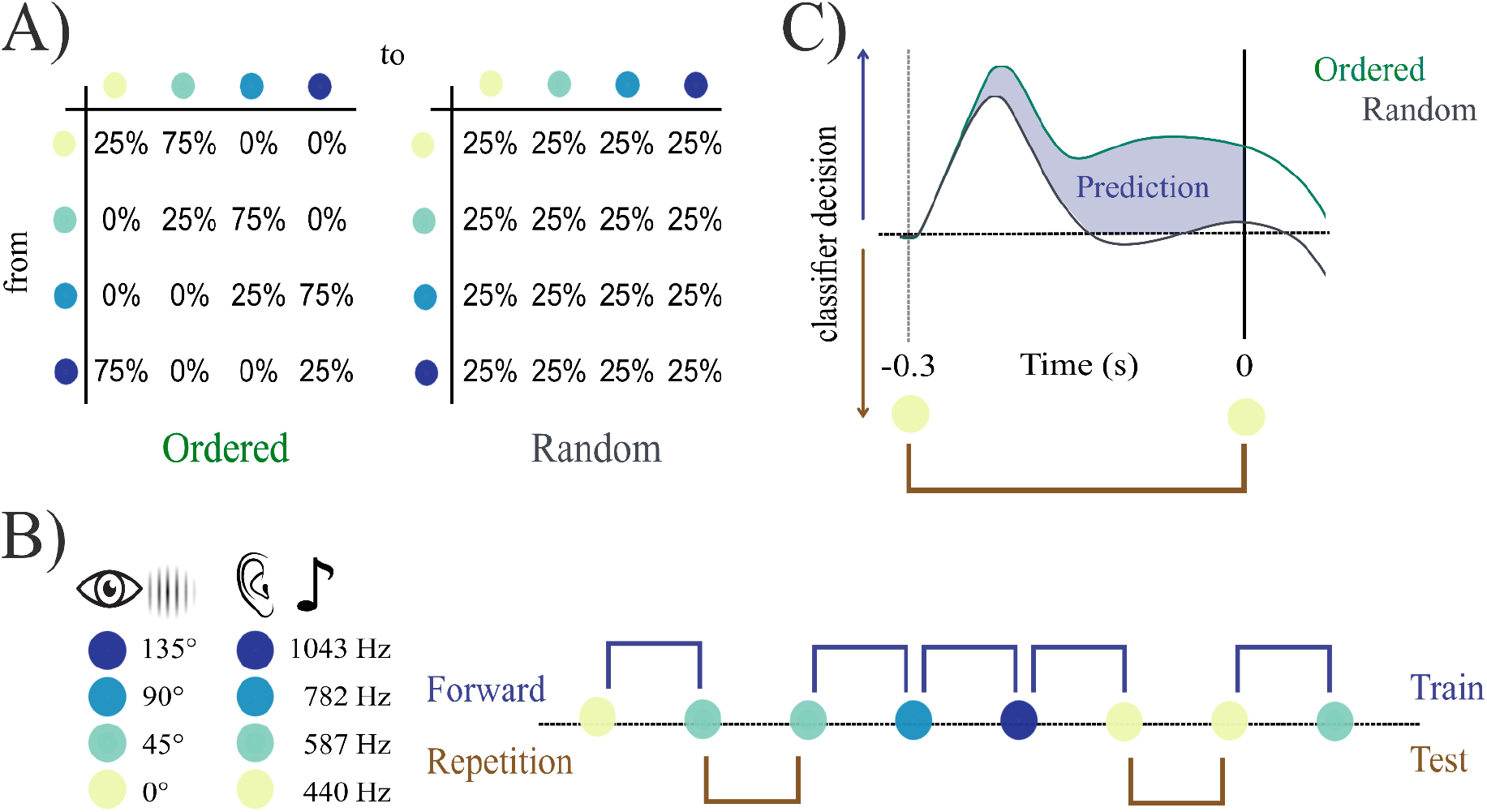
Quantification of individual prediction tendency. **A)** Participants were presented with sequences of four different stimuli. Transitional probabilities varied according to two entropy conditions (ordered vs. random). **B)** Visual stimulation consisted of gabor patches in four different orientations, auditory stimulation consisted of pure tones at four different frequencies. An LDA classifier was used to decode stimulus feature from brain activity across time, trained on (high-probability) ordered forward transition trials and tested on all repetition trials. **C)** Expected classifier decision values contrasting the brain’s pre-stimulus tendency to predict a forward transition between the two entropy levels. Individual prediction tendency quantification results from the summed difference between conditions (ordered > random) over pre-stimulus time.

### MEG Data Acquisition and Preprocessing

A whole-head MEG system (Elekta Neuromag Triux, Elekta Oy, Finland), placed within a standard, passive magnetically shielded room (AK3b, Vacuumschmelze, Germany), was used to capture magnetic brain activity with a sampling frequency of 1 kHz (hardware filters: 0.1 - 330 Hz). The signal was recorded with 102 magnetometers and 204 orthogonally placed planar gradiometers at 102 different positions. In a first step, a signal space separation algorithm, implemented in the Maxfilter program (version 2.2.15) provided by the MEG manufacturer, was used to clean the data of external noise and realign data from different blocks to a common standard head position. Data preprocessing was performed using Matlab R2020b (The MathWorks, Natick, Massachusetts, USA) and the FieldTrip Toolbox (Oostenveld et al., 2010). To identify eye blinks and heartbeat artifacts, 50 independent components were identified from filtered (0.1 - 100 Hz) continuous data of the first experimental block of both modalities (auditory + visual). On average, 2.6 (range = 2 - 5) components were removed for each subject. All data was filtered between 0.1 Hz and 30 Hz (Kaiser windowed finite impulse response filter) and downsampled to 100 Hz. Then, the data of each block was epoched into segments of 1200 ms (from 400 ms before stimulus onset to 800 ms after onset) for further analysis (as in Schubert et al., 2023).

### Decoding Analysis

Multivariate pattern analyses were carried out using the MVPA-Light package (Treder, 2020) and were conducted separately for each modality. Prior to the classification analysis, we excluded the first 20 trials after each new entropy onset (resulting in a total of 2720 trials per entropy and modality). This decision was based on the number of trials, it would take an ideal Bayesian observer to reach above 0.5 posterior probability for a certain stimulus class in an ordered context, which was modeled using the HGF toolbox (Frässle et al., 2021). A multi-class linear discriminant analyser (LDA) was used to classify stimulus feature (i.e. 1 - 4, sound frequency or gabor-patch orientation) from brain activity between -0.3 and 0.3 ms in a time-resolved manner. Based on resulting classifier decision values (i.e. d1 - d4 for every test-trial and time-point), we calculated the individual prediction tendency within each modality (as in Schubert et al., 2023): We define individual prediction tendency as the tendency to pre-activate sound frequencies of high probability (i.e. a forward transition from one stimulus to another: 1→2, 2→3, 3→4, 4→1). In order to capture any prediction-related neural activity, we trained the classifier exclusively on ordered forward trials. Afterwards, the classifier was tested on self-repetition trials, providing classifier decision values for every stimulus feature, which were then transformed into corresponding transitions (e.g. d1(t) | 1(t-1) “dval for 1 at trial t, given that 1 was presented at trial t-1” → repetition, d2(t) | 1(t-1) → forward,…). The tendency to represent a forward vs. repetition transition was contrasted for both ordered and random trials. Using self-repetition trials for testing, we ensured a fair comparison between the ordered and random contexts (with an equal probability and the same preceding bottom-up input) and that test trials were always different from training trials. To ensure that classifier decision could also not have been biased by the stimulus presentation at t-2, ordered and random trials were matched for the preceding stimuli at t-1 (using only repetitions from t-1→ t) and t-2 (using only forward transitions from t-2 → t-1). This resulted in a total of 84 - 124 (mean = 108.6) test trials in the auditory modality and 92 - 126 (mean = 108.77) test trials in the visual modality per subject and entropy condition (ordered trials were randomly subselected to match the number of random trials). Thus, our time window of interest (−0.3 s - 0 s) should not contain any carry-over effects of preceding stimuli (note that stimulus identity can be classified until 700 ms after presentation, see **Fig. S1, Appendix**). Additionally, we also conducted a similar analysis (which can be found in more detail in Demarchi et al., 2019), where we trained the classifier on random trials and time-generalised its performance to capture potential prestimulus representations in both conditions (see **Appendix**). Based on the consideration that stimulus “predictions” are not necessarily the same as poststimulus representations, we focus our interpretation on the analysis approach in which the classifier has been trained on ordered trials (and was able to capture “prediction”-specific patterns).

Thus, we quantified “prediction tendency” as the classifier’s pre-stimulus tendency to a forward transition in an ordered context exceeding the same tendency in a random context (which can be attributed to carry-over processing of the preceding stimulus). Then, using the summed difference across pre-stimulus times, one value can be extracted per subject, per modality.

### Statistical Analysis

The classifier’s tendency toward a forward transition in an ordered context was compared to the same tendency in a random context, using a cluster-based permutation t-test (using 10,000 random permutations, a cluster-alpha of 0.025 and a monte-carlo critical alpha of 0.025) over a time window from -0.3 s to 1.5 s. This contrast was calculated for both modalities separately. Afterwards, the summed differences across pre-stimulus time was extracted for each subject and modality, resulting in individual prediction tendency values. These single values were then used to calculate the pearson’s correlation coefficient between auditory and visual prediction tendency. To extend our findings by classical estimates of internal consistency, we also calculated Cronbach’s alpha and the Spearman-Brown reliability estimates, assuming that prediction tendency can be understood as a multi-dimensional concept, here captured with a two-item (auditory, visual) scale. Additionally, we also calculated the reliability of predictions between modalities (Spearman-Brown’s cofficient: rho = 2*r/(1+r); formula taken from Eisinga et al., 2013) for all possible pre-stimulus time-by-time combinations. All analyses were done in MATLAB version R2021a.

## RESULTS

### Anticipatory predictions can be found mainly in the auditory modality

To investigate anticipatory prediction, participants were presented a sequence of four different (either visual or auditory) stimuli, which changed according to predefined transitional probabilities of two entropy levels: ordered vs. random. In a first step, we were interested if we would see a pre-activation of high-probability stimulus features in brain activity across subjects. Cluster-based permutation showed an overall auditory prediction tendency (i.e. ordered > random) in two pre-stimulus clusters from -0.23 s to -0.2 s (p = 0.035) and from -0.17 s to -0.07 s (p = 6.9*10^-4^). For visual prediction tendency, we found only a trend suggesting prestimulus predictions in a short cluster from -0.24 s to -0.22 s (p = 0.056). Using a different approach, where we trained on random trials and time-generalised classifier performance into a prestimulus window we found a similar picture indicating stimulus pre-activations in the auditory but not in the visual modality (see **Fig. S2, Appendix**). These results indicate that, irrespective of individual differences, there seems to be an overall tendency to anticipate stimulus features of increased probability in the auditory modality (see **Fig. 2A**). In the visual modality, however, the results are less conclusive, suggesting that anticipatory predictions might not be a general group phenomenon.

**Fig. 2:**
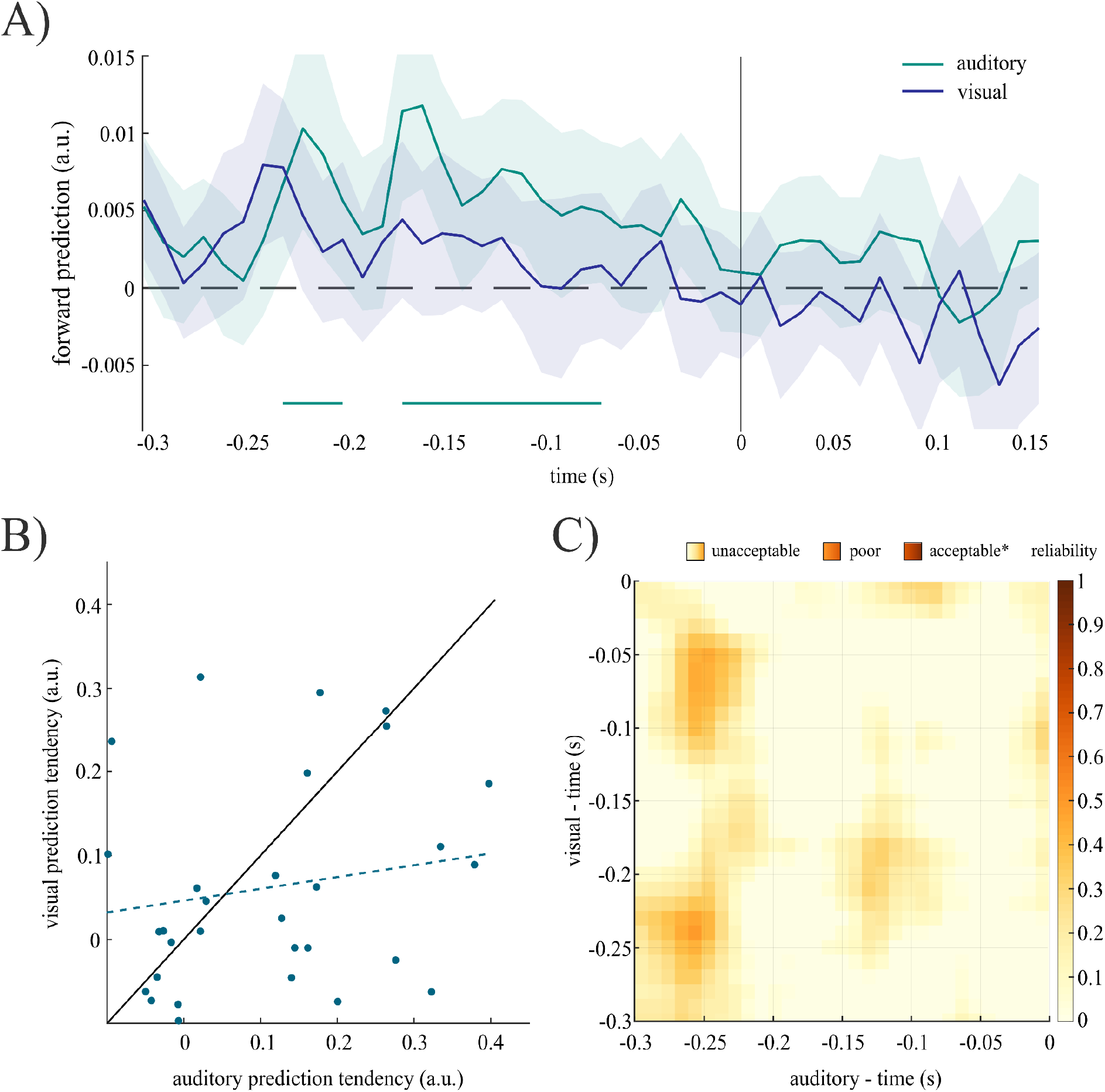
**A)** Time-resolved prediction tendency for auditory and visual modalities: On a group-level there seems to be an anticipatory tendency to represent stimulus features of high probability (i.e. forward transition) in a predictable context in the auditory modality, but not in the visual modality (y-axis represents the difference in dvals (ordered - random) for a forward transition; solid lines on x-axis indicate significant time-points (based on a random permutation test using an alpha level of 0.025)). **B)** Correlation of individual prediction tendency between modalities: There is no significant correlation between auditory and visual prediction tendency (the solid line represents a perfect generalisation from one modality to another; the dashed line indicates the least-squared regression line). **C)** Temporal generalisation of forward predictions across modalities: early anticipatory predictions in the auditory modality seem to generalise best to late anticipatory predictions in the visual modality, however only poor reliability is indicated overall (colormap represents Spearman-Browns coefficient, * taken from Tavakol & Dennick, 2011; N = 35).

### Individual prediction tendency is not correlated between modalities

In a second step, we wanted to further investigate prediction tendency as a subject-specific, potentially unified concept. For this purpose, we looked particularly into the generalisation between auditory and visual modalities. Using the summed prediction tendency values across pre-stimulus time (−0.3 s - 0 s), one value was extracted per subject, per modality. Even though individual prediction tendency values were, on average, higher for the auditory (mean = 0.076, sd = 0.164) than the visual (mean = 0.059, sd = 0.127) modality, there was no significant difference in a one-sample t-test (t = 0.533, 95% CI = [−0.048, 0.082]). In total, 20 subjects showed a stronger auditory prediction tendency, compared to 15 subjects with stronger prediction tendency in the visual domain (see **Fig. 2B**). Crucially, we found no relationship of individual prediction tendency between modalities, using pearson’s correlation (r = 0.18, p = 0.300; see **Fig. 2B**). To test the assumption that both measures represent the same concept of individual prediction tendency, we further calculated Cronbach’s alpha and Spearman-Brown’s rho to estimate internal consistency (see Table 1). In order to be able to interpret the reliability across modalities we also calculated the internal (split-half) consistency within each modality. We found a significant positive correlation between prediction tendency estimated from one half (trials were randomly selected) and the other half in the auditory (r = 0.74, p = 3.73*10^-7^) as well as in the visual modality (r = 0.58, p = 3.00*10^-4^). Because there seems to be some disagreement as to which reliablity statistic should be used for measures comprising two items, we decided to report all three here (Eisinga et al., 2013). In an attempt to account for individual temporal dynamics of each modality, we also estimated the reliability for all possible pre-stimulus time-by-time combinations. It seems that early auditory predictions generalise best to late visual predictions, with overall low reliability (see **Fig. 2C**). Crucially, based on guidelines from classical test theory, our results indicate that the effect is too weak to provide a reliable generalisation from one modality to another (Tavakol & Dennick, 2011). In sum, these findings suggest that individual prediction tendencies don’t necessarily generalise across different modalities.

**Table 1:** Comparison of correlation coefficient, cronbach alpha and spearman brown coefficient

|  | Pearson's r | Cronbach's alpha | Spearman-Browns rho |
| --- | --- | --- | --- |
| Reliability across modalities | 0.18 | 0.298 | 0.306 |
| Auditory (split-half) reliability: | 0.74 | 0.850 | 0.851 |
| Visual (split-half) reliability: | 0.58 | 0.710 | 0.730 |

Note: Reliability was calculated treating *prediction tendency* as a two-item (*auditory, visual*) scale.

## DISCUSSION

Predictive processing has been found throughout the range of human cognition: in language perception and production (e.g. Pickering & Garrod, 2013), object recognition (e.g. Oliva & Torralba, 2007), motor perception and action (e.g. Shipp et al., 2013), emotional awareness (e.g. Smith et al., 2019), and decision making (e.g. Summerfield & de Lange, 2014). However, the extent to which all these predictions can be attributed to a collective mechanism remains unclear. Beyond that, several conceptual and mathematical predictive brain models have been proposed to explain cognitive and perceptual pathologies in clinical neuroscience (for a review see Smith et al., 2021). Considering that psychiatric conditions such as depression, autism or schizophrenia are assumed to be highly individualized and stable across time and situations, estimations of prediction tendency are likely to be interpreted as a “trait-dependent” rather than a “state-dependent” measure. In this regard, prediction tendency should a) vary considerably across individuals, b) show long-term stability within an individual and c) generalise across different situations. Yet research that validates this claim is scarce. In the current study, we aimed to investigate to what extent the individual tendency to anticipate sensory features of high probability generalises across modalities. Even though aberrant predictive tendencies, especially linked to psychosis, have been proposed for auditory (Corlett et al., 2019) as well as visual perception (Adams et al., 2012), this is, to our knowledge, the first study to directly compare individual prediction tendencies between two modalities. Our results show that under conditions of low entropy, stimulus features of high probability are preactivated in the auditory modality irrespective of individual differences. In the visual modality, however, evidence for such preactivations is less clear. Crucially, this tendency to predict sensory events seems not to correlate between the two modalities, suggesting that the quantification and interpretation of prediction tendencies cannot be generalised.

Even though we only found inconclusive evidence for anticipatory predictions in the visual modality on a group-level (with the lowest p-value of 0.056 not reaching significance), we argue that our focus lies on individual differences, which should be compared across modalities independent of their group-level significance. However, we can conclude that less than 4% of total variance in visual prediction tendency can be explained by auditory prediction tendency and vice versa. We argue that this seems too weak to support the assumption that individual prediction tendency can be generalised from one modality to another. Considering that the same metric was applied in both cases, we orient ourselves primarily on standards that are also used to assess reliability, for which acceptable statistics usually range between 0.7 and 0.95 (Tavakol & Dennick, 2011). Additionally, it should be noted that our prediction tendency quantification was derived from the decoding of stimulus features from brain activity, a measure that is highly affected by a subject-specific signal-to-noise ratio (as a result of anatomical differences or individual distributive properties, a problem often discussed when calculating source localisation; Puce & Hämäläinen, 2017). It can therefore be assumed that decoding accuracies are correlated within subjects, and it should be considered that a certain proportion of the relationship is driven by individual differences in signal-to-noise ratio alone. In addition, it should be considered that our quantification might not be a “pure” measure of prediction tendency, which reduces internal consistency, even within a modality. However, reliability statistics indicate acceptable internal consistency within each modality (see Table 1). In sum, we support the notion that scientific evidence should not be judged solely on the basis of its significance, but that the strength of effects should be evaluated within the framework of underlying assumptions. We therefore conclude that our estimates of reliability are too weak to allow a generalisation across modalities.

Our results indicate that individuals who show a stronger auditory prediction tendency don’t necessarily show a similarly strong tendency for visual predictions. Indeed it seems that overall individuals don’t show similar anticipatory predictions in the visual modality as they do in the auditory modality. There are different possible explanations for these findings. In the current paradigm, stimulus predictability was modulated via transitional probabilities in a sequence of events, a feature that is quite characteristic for auditory stimuli, such as music and speech, but maybe less so for visual stimulation (ten Oever et al., 2014). To make optimal use of top-down inferences in auditory processing, sensory information has to be integrated over (past) time (and extrapolated into the future) and, crucially, has to be accurately located within time. Although visual processing has to meet similar requirements, particularly when confronted with moving objects, the motivation for anticipatory representations might be different (e.g. to facilitate suitable motor action in time), potentially resulting in increasingly goal-driven predictions in the visual modality (for a review see Fiehler et al., 2019). Such differences in motivation could very well lead to modality-specific implementations of anticipatory predictions, which might explain the weak generalisation between modalities in our results.

Indeed, it seems that the extent to which an observer (versus a listener) has to be actively engaged in the sampling of information is different between vision and audition. For example, information that gradually unfolds over time, such as a spoken sentence, can (under optimal noise conditions) be received without any active adjustments from the listener (although see, for example, Gehmacher et al., 2022, for top-down influence on auditory nerve activity). Brain responses to unexpected changes in auditory stimulation (the classic “mismatch negativity”), for example, have even been found during sleep (Loewy et al., 1996). Meanwhile visual information, which is usually spread throughout space, extending the foveal field of the observer, requires active effort (e.g. eye, head or body movements) to be sampled. According to some theories, however, active sampling of sensory evidence is crucial for the formation of internal models and the predictions that come from them (Friston et al., 2017). Since our paradigm was entirely passive, no beneficial gain was to be expected from focusing on the sequences. Thus, participants likely did not allocate their attentional resources to them. If the proposed link between predictions and active engagement is stronger in vision, a lack of attention and goal-directedness could explain why we found automatic anticipatory predictions in the auditory but not in the visual modality. We suggest that future research should investigate the (potentially different) roles of attention for predictive processing in vision versus audition.

A different explanation is that statistical learning (SL), which has been defined as “a general capacity for picking up regularities”, is qualitatively different between modalities (Siegelman et al., 2017). This individual capacity is implicit in our conceptualisation of “prediction tendency”, as we quantify to what extent an individual anticipates sensory features of high probability (compared to a low probability context). Our results are consistent with findings in that area, which suggest that SL can also not be generalised across different modalities (Siegelman et al., 2017). Crucially, it has been found that visual SL performance is higher (comparable to auditory SL performance for sequences) when information is distributed spatially, compared to when visual input was distributed temporally (Conway & Christiansen, 2009). This finding supports our interpretation that overall prediction tendency for sequential input is different between the auditory and the visual modality. Finally, it should be noted that in sequential SL paradigms, the observers internal model of input probability is based on past experience of a few trials. Therefore, individual prediction tendency might not be stationary but fluctuate even within a given regularity context (see for example Notaro et al., 2019). A related limitation of the current paradigm is that our quantification of prediction tendency includes the observers ability to distinguish between an ordered and a random context. This way, strong priors that are immune to changes in contextual probability (i.e. anticipatory predictions towards a favoured transition in a random context) remain undetected. An interesting future perspective would be to look into the generalisability of predictive tendencies across different regularity contexts.

### Conclusions and implications for future research

Predictive processing theories, which model the brain as a “prediction machine”, are able to explain a wide range of cognitive functions, including learning, perception and action. Furthermore, it seems to be widely accepted that aberrant prediction tendencies play a crucial role in long-term psychiatric disorders. Crucially, in clinical neuroscience, prediction tendency is often implicitly conceptualized as an individual trait or as a tendency that generalises across situations, yet research to validate this claim is scarce. In the current study, we found evidence for anticipatory predictions in the auditory but not in the visual modality. Furthermore, our results suggest that quantification and interpretation of individual prediction tendency cannot be generalised across modalities. This emphasises the importance of research, further investigating and validating the implicit assumption of “prediction tendency” as a trait, since this assumption bears concrete implications for the growing field of computational modeling in neuroscience and psychiatry. We propose that future research focuses on the validation of state- vs. trait-dependent assumptions in predictive processing theories. Furthermore, we strongly suggest investigation of the intraindividual stability of prediction tendency within and across different modalities.

## Acknowledgements

J.S. is supported by the Austrian Science Fund (FWF; Doctoral College “Imaging the Mind”; W 1233-B). N.S. i supported by the Austrian Science Fund, P31230 (“Audiovisual speech entrainment in deafness”) and P34237 (“Impact of face masks on speech comprehension”). Thanks to the whole research team. Special thanks to Manfred Seifter for his support in conducting the measurements.

## Author Contributions

J.S. designed the experiment, analyzed the data, generated the figures, and wrote the manuscript. N.S. recruited participants, supported the data analysis, and edited the manuscript. N.W. designed the experiment, acquired the funding, supervised the project, and edited the manuscript.

## APPENDIX

## Poststimulus decoding of auditory and visual features

In order to generally verify feature decodability from neural activity and to investigate the duration for which a given stimulus representation remains sustained in both modalities, we conducted a classification analysis that focused solely on trials in the random condition. We trained and tested an LDA classifier in a time-resolved manner from -0.2 s to 1 s using a 5-fold cross validation approach. We find that auditory features (i.e. sound frequencies) can be classified from brain activity starting 0.06 s until 0.67 s following sound onset. Similarly visual features (i.e. gabor patch orientation) can be classified from 0.07 s to 0.74 s after stimulus onset. (Note that in this case “decodability” was defined as the lower boundary of a 95%CI exceeding the chance level of 0.25, which was not corrected for multiple comparisons, since we wanted to gain a liberal estimate for the duration of feature-specific information.) We conclude that auditory as well as visual features can be classified from brain activity starting at ∼100 ms until ∼700 ms after stimulus onset. Therefore, we argue that 700ms is the time-window in which potential carry-over effects should be controlled for when comparing classifier performance between an ordered (systematic stimulus variation) and a random (unsystematic variation) context.

**Fig. S1:**
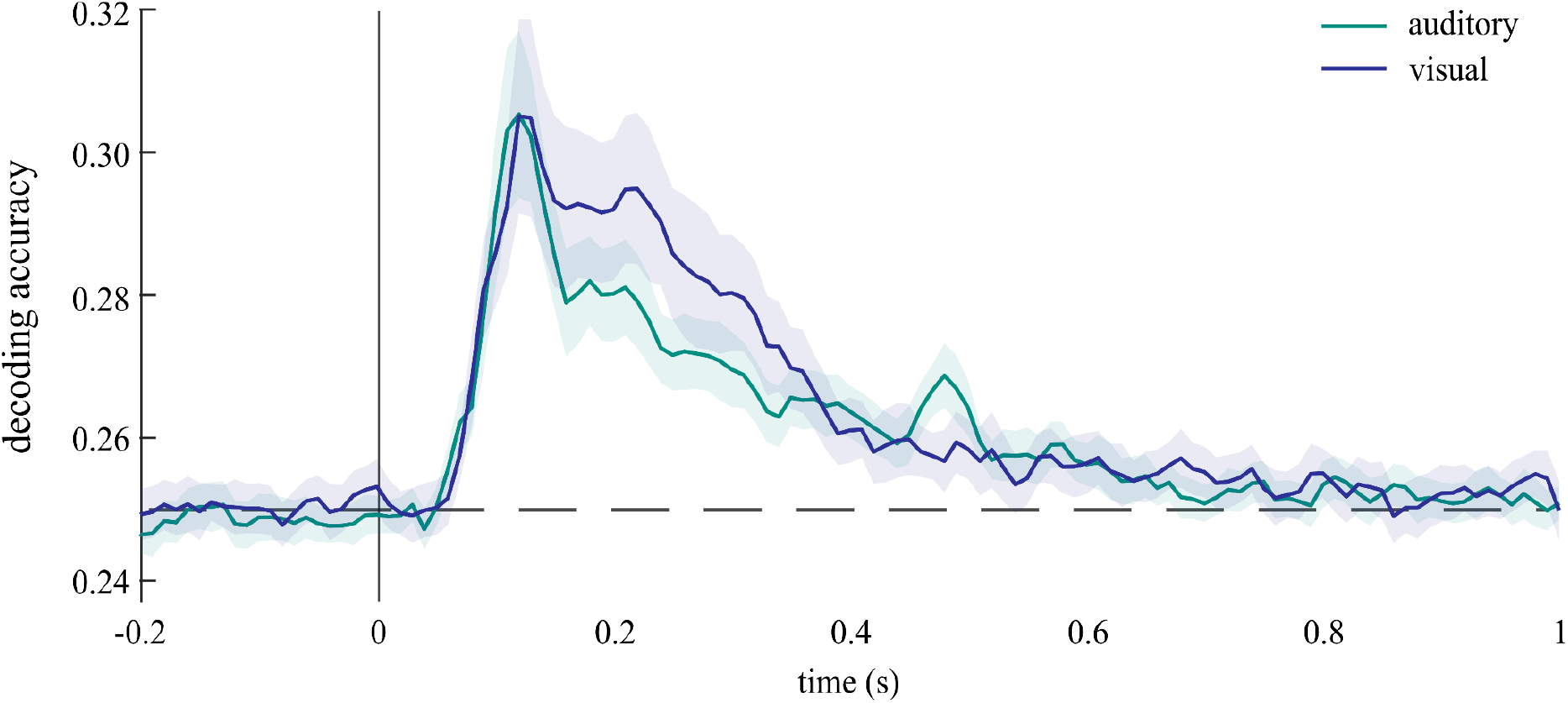
Time-resolved decoding accuracy for auditory and visual features: sound frequency as well as gabor patch orientation can be classified from brain activity from ∼100 ms until ∼700ms after stimulus onset in a random context (shaded area indicates 95%CI, dashed line shows the chance level of 0.25; N = 35).

## Anticipatory predictions can be found mainly in the auditory modality

To quantify “prediction tendency”, we compared the classifier’s prestimulus tendency for a highly probable forward transition in an ordered context to the same tendency in a random context. Cluster-based permutation showed an overall auditory prediction tendency (i.e. ordered > random) in two prestimulus clusters from -0.23 s to -0.2 s (p = 0.035) and from -0.17 s to -0.07 s (p = 6.9*10-4). For visual prediction tendency, we found only a trend suggesting prestimulus predictions in a short cluster from -0.24 s to -0.22 s (p = 0.056).

**Fig. S2:**
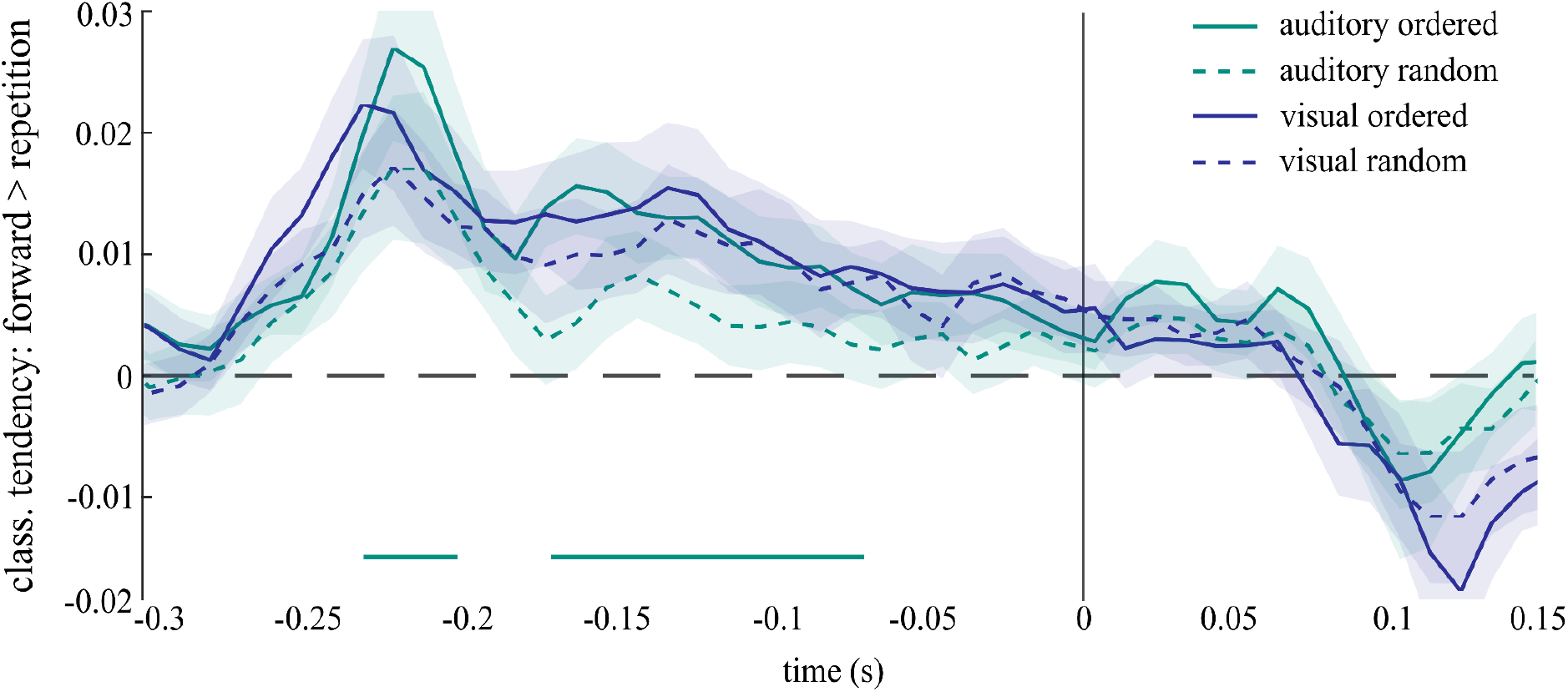
Tendency to represent a “forward” transition (compared to a “repetition” transition), separately for different entropy levels and modalities. On a group-level there seems to be an anticipatory tendency to represent stimulus features of high probability (i.e. forward transition) in a predictable context in the auditory modality, but not in the visual modality. (Note that this Figure shows the same data as **Fig. 2A**, but with separate lines for each entropy condition. Y-axis represents the classifier dvals for a “forward” transition before subtraction (ordered - random) and the solid lines on x-axis indicate significant time-points (ordered > random); N =35).

Additionally, we conducted a second analysis (see also Demarchi et al., 2019), where we trained the classifier on poststimulus brain activity of random trials and time-generalised its performance to capture potential prestimulus representations in ordered as well as random trials (for more information on the temporal generalisation approach see also King & Dehaene, 2014). One advantage of this approach is that, as the classifier is trained on random trials only, it remains unbiased and does not require for the preceding stimuli to be matched between conditions. Using a cluster-based permutation test, we compared the classifiers performance to generalise from post-into prestimulus intervals between the ordered and the random condition separately for each modality. We found that in the auditory modality, there was a positive prestimulus cluster indicating that, in an ordered context, brain activity patterns preceding a stimulus were more similar to pure bottom-up processing patterns than in a random context (see **Fig. S3**). We conclude that anticipatory predictions contain feature-specific activations similar to those following sound onset. In the visual modality, however, we found no such cluster (see **Fig. S3**). In sum, these findings suggest that anticipatory predictions can be found mainly, if not exclusively, in the auditory modality.

**Fig. S3:**
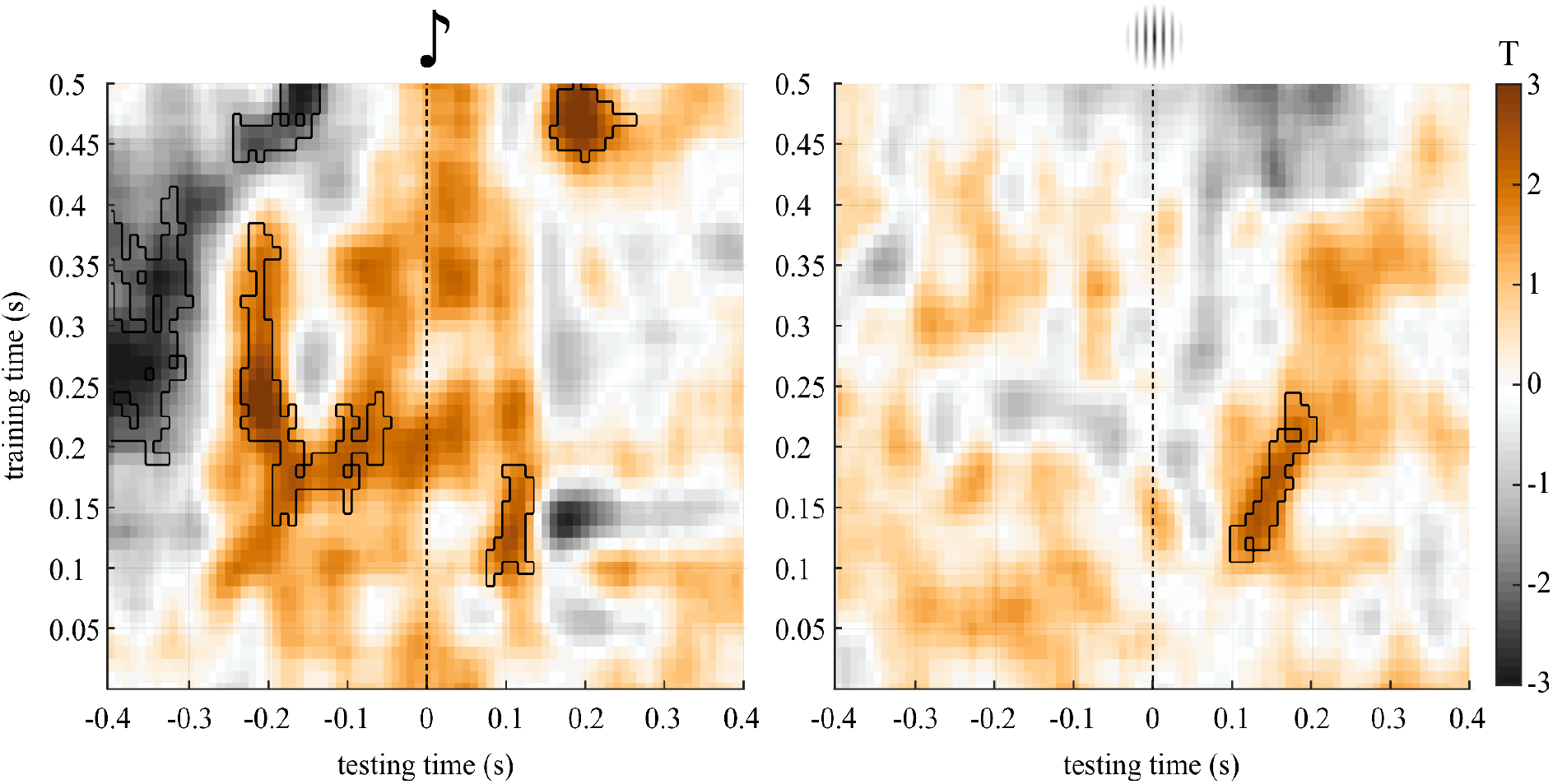
Temporal generalisation of feature-specific activations in the auditory (left) and visual (right) modality. Left: In the auditory modality there is a significant difference in the generalisation from post- to prestimulus processing between entropy levels. This suggests that in an ordered, but not in a random, context people generate feature-specific anticipatory predictions, that resemble bottom-up processing. Right: In the visual modality, however, we find no significant evidence for a regularity-dependent generalisation from post- to prestimulus acitvations. (Y-axis represents classifier poststimulus training-time and X-axis represents classifier pre- and poststimulus testing-time; T-values are shown in color and marked outlines indicate a significant cluster in the comparison ordered vs. random; N = 35).

